# Unsupervised explainable AI reveals similar oligonucleotide-usage zones matching the highest-resolution human chromosome bands

**DOI:** 10.64898/2026.03.09.710446

**Authors:** Toshimichi Ikemura, Yuki Iwasaki, Kennosuke Wada, Yoshiko Wada, Takashi Abe

**Affiliations:** Faculty of Bioscience, Nagahama Institute of Bio-Science and Technology, Tamura-cho 1266, Nagahama, Shiga 526-0829, Japan; Association for Propagation of the Knowledge of Genetics, National Institute of Genetics, Yata 1111, Mishima, Shizuoka 411-8540, Japan; Graduate School of Science and Technology Electrical and Information Engineering, Niigata University, Niigata 950-2181, Japan

**Author notes:** Corresponding authors. (T.I.).

**Keywords:** artificial intelligence, machine learning, oligonucleotide odds ratio, Self-Organizing Map, epigenomics, prophase 2000 bands

## Abstract

Unsupervised and explainable AI can uncover genomic features that extend beyond human expectations. Oligonucleotides such as penta- and hexanucleotides often function as core binding motifs for regulatory proteins, and their usage provides a powerful tool in functional genomics. We applied an unsupervised, explainable AI approach to odds ratio (observed/expected) profiles of all 1-Mb euchromatic fragments in the human genome. This odds-ratio analysis identified oligonucleotide features independent of mononucleotide composition, thereby highlighting functional roles of oligonucleotide motifs. AI-based clustering of all 1-Mb euchromatic fragments, using either penta- or hexanucleotide odds ratios (1,024 or 4,096 variables), unexpectedly revealed nearly 2,000 distinct zones, despite the large differences in dimensionality. If these ∼2,000 zones represent biologically meaningful segmentations, comparable structures would be expected to emerge when other oligonucleotide types are analyzed. Consistent with this expectation, CG-containing penta- and hexanucleotides (244 and 1,185 variables) produced comparable ∼2,000 zones, indicating that the underlying segmental structures reflect fundamental functional divisions within the genome. Human chromosomes exhibit well-established Giemsa-banding patterns comprising 850 bands at prometaphase and 2,000 bands at prophase. Since single-nucleotide coordinates are available for the 850 bands, we identified a diagnostic oligonucleotide set that distinguishes Giemsa-negative and -positive regions. Computational pseudo-band reconstruction based on this set generated genome segmentations that more closely paralleled the AI-derived ∼2,000 clusters than the 850 bands. These unexpected findings indicate that AI captures the characteristic features of chromosomal bands and can predict high-resolution banding from genome sequences alone, thereby bridging classical cytogenetics and modern AI-based genomics.

## INTRODUCTION

The oligonucleotide compositions of diverse species have been characterized (Nussinov 1984; Kryukov et al. 2012), and interspecies variation in short oligonucleotide composition has been referred to the genomic signature (Karlin et al. 1997). The Batch-Learning Self-Organizing Map (BLSOM), an unsupervised machine-learning method, for oligonucleotide analysis has enabled robust species-dependent clustering of genomic fragments (Abe et al. 2003, 2005). Applied to vertebrate genomes, BLSOM also uncovered Mb-scale intragenomic separation defined by distinctive oligonucleotide compositions (Abe et al. 2006; Katsura et al. 2021). Moreover, analysis of the hg19 and hg38 assemblies of the human genome revealed the chromosome-dependent separation of centromeric and pericentromeric constitutive heterochromatin sequences, which contain dense clusters of repetitive sequences harboring core motifs for transcription-factor binding (Iwasaki et al. 2013, 2022; Wada et al. 2015, 2020). The present study applies BLSOM to characterize Mb□scale oligonucleotide composition primarily in human euchromatic regions. For clarity, the following abbreviations are used for mono- and oligonucleotides of varying lengths hereafter: Oligo (general), Mono, Di, Tri, Tetra, Penta, and Hexa. Plurals are denoted by adding an “s.” Oligo sequences carry biological significance. For example, the CG dinucleotide is a major site of cytosine methylation involved in epigenomic regulation (Kim and Costello 2017), and longer Oligos often serve as regulatory core motifs, including transcription-factor binding sites. The segmental profile of Oligo composition across chromosomes thus provides a powerful framework for investigating the functional organization of the human genome.

Oligo composition is inevitably influenced by Mono composition, particularly G+C%. Bernardi and colleagues (Macaya et al. 1976; Bernardi et al. 1985) demonstrated that warm□blooded vertebrate genomes are partitioned into large segmental structures (>300 kb) defined by G+C%, termed isochores. Isochores correlate with various genomic features, including gene density, replication timing, nuclear organization, and cytogenetic banding; in the human genome, ∼3,200 isochores have been delineated (Costantini and Bernardi 2008; Bernardi 2009). Our group has also characterized large G+C%-based segmental structures, first noted through codon-usage analyses (Ikemura 1985; Aota and Ikemura 1986), and has shown that these segments relate to chromosome bands (Ikemura and Aota 1988; Ikemura et al. 1990; Ikemura and Wada 1991). The present study focuses on the odds ratio (observed/expected) to characterize Oligo-composition profile independent of Mono composition: “observed” refers to the measured frequency, whereas “expected” refers to the frequency predicted from Mono composition. Oligo odds ratio (OligoOdds) shifts the focus from Mono bias to intrinsic Oligo features that are likely linked to biological functions. Penta- and HexaOdds BLSOM analyses of the T2T genome (Aganezov et al. 2022) reveal chromosome□dependent self-organization of 1□Mb euchromatic sequences, partitioning euchromatin into ∼2,000 discrete zones. We assess their biological relevance by comparing these zones with chromosome bands as follows.

Chromosome banding techniques have been established across higher eukaryotes, and despite differences in staining principles, they consistently yield comparable banding patterns (Holmquist 1992). These patterns reflect fundamental genome features, including segmental G+C% organization, chromatin compaction, replication timing, gene density, and nuclear spatial organization (Holmquist 1992; Craig and Bickmore 1993; Costantini and Bernardi 2008). Band patterns are therefore regarded as indicators of functional genome segmentation. FISH mapping of numerous BAC clones enabled estimation of the genomic coordinates of 850 Giemsa□stained prometaphase bands (Cheung et al. 2001). The UCSC Genome Browser (Perez et al. 2025) defined their boundary coordinates on the T2T genome at single□base resolution for computational use, yielding an average band length of 3.65 Mb. Although anchoring these Mb□scale bands to genomic coordinates enables nucleotide□level analyses, positional uncertainty inherent to BAC□FISH mapping persists. Notably, another Giemsa staining pattern comprising 2000 bands in prophase represents the highest-resolution banding for humans (Yunis 1981), and narrow sub□bands at this level are typically nested within the broader structures of 850 bands. Using positional information for 850 bands, we identified an OligoOdds set that distinguishes Giemsa-negative and -positive bands. Applying this diagnostic set to computational pseudo-band reconstruction produces segmentation patterns closely matching the ∼2,000 AI-defined zones, despite being anchored to the 850□band framework.

## RESULTS

### BLSOM Analysis Across the Entire T2T Sequence

We applied BLSOM to analyze Di to Penta compositions across the complete human T2T genome. Results for Di to Tetra compositions are provided in Supplementary Fig. S1. Figure 1A shows Penta composition of 1-Mb fragments. Fragmentation was performed using a 1-Mb sliding window started at the short arm terminus of all chromosomes and shifted with a 10□kb step. As the method relies on unsupervised machine learning, no chromosomal labels were provided during the learning process. Thus, clustering (self-organization) emerged solely from Oligo-composition similarity. For this and subsequent analyses, the map size is adjusted so that each node contains an average of ten sequences. Nodes containing fragments from a single chromosome are colored, while nodes with mixed origins are shown in black.

**Figure 1.**
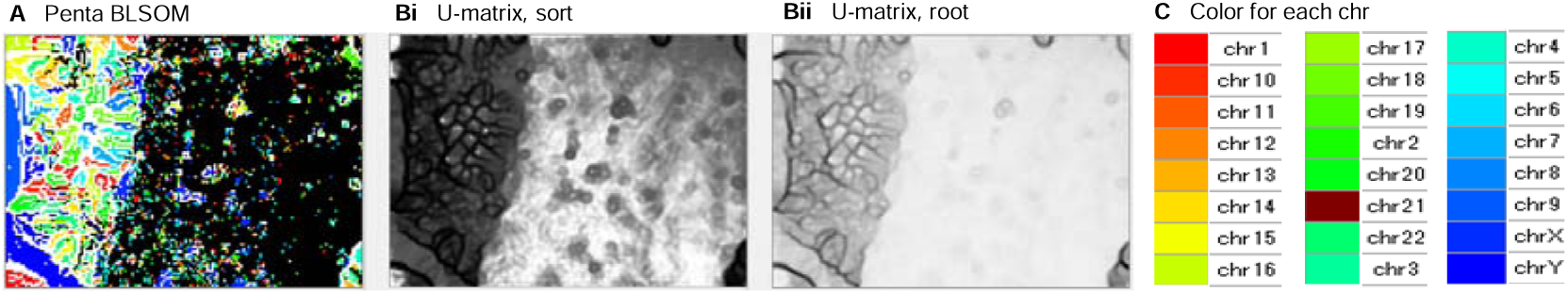
BLSOM for Penta composition of 1-Mb windows across the T2T genome. (**A**) Nodes containing sequences from multiple chromosomes are shown in black; those containing sequences from a single chromosome are colored. (**Bi, Bii**) U-matrix showing Euclidean distances between neighboring nodes. Shading reflects either the sum of vector distances to nearest neighbors (**Bi**) or its square root (Bii). (**C**) Chromosome and color correspondence. In this study, the specific color assigned to each chromosome is not essential. However, to distinguish chr21 from chr22 in subsequent figures, the color of chr21 was adjusted to improve its visual distinction.

The left third of the map is dominated by colored nodes, reflecting chromosome-dependent clustering consistent with our previous study for hg38 (Wada et al. 2020); these clusters correspond to centromeric and pericentromeric constitutive heterochromatin, as noted in the Introduction. Figure 1B displays U□matrix patterns (Ultsch 1990) representing Euclidean distances between neighboring nodes in the BLSOM shown in Fig. 1A. Node shading denotes either the sum of vector distances to nearest neighbors (Fig. 1Bi) or its square root (Fig 1Bii), which distinguishes between local and global distances. In the leftmost third of the map (Fig. 1A), which corresponds to the constitutive heterochromatin as mentioned above, chromosome□dependent separations align with distinct black boundaries in the U-matrix (Fig. 1B), revealing inter□chromosomal differences in Penta composition.

### BLSOM of Di Composition and Odds Ratio in Euchromatin

We previously characterized Oligo usage in centromeric and pericentromeric heterochromatin in detail (Iwasaki et al. 2013, 2022; Wada et al. 2015, 2020). Here, BLSOM analysis is extended to euchromatic regions, which remain less explored. Using the same procedure as described in Fig. 1, we analyzed the euchromatic regions with a 1-Mb sliding window and a 10-kb step (Fig. 2).

**Figure 2.**
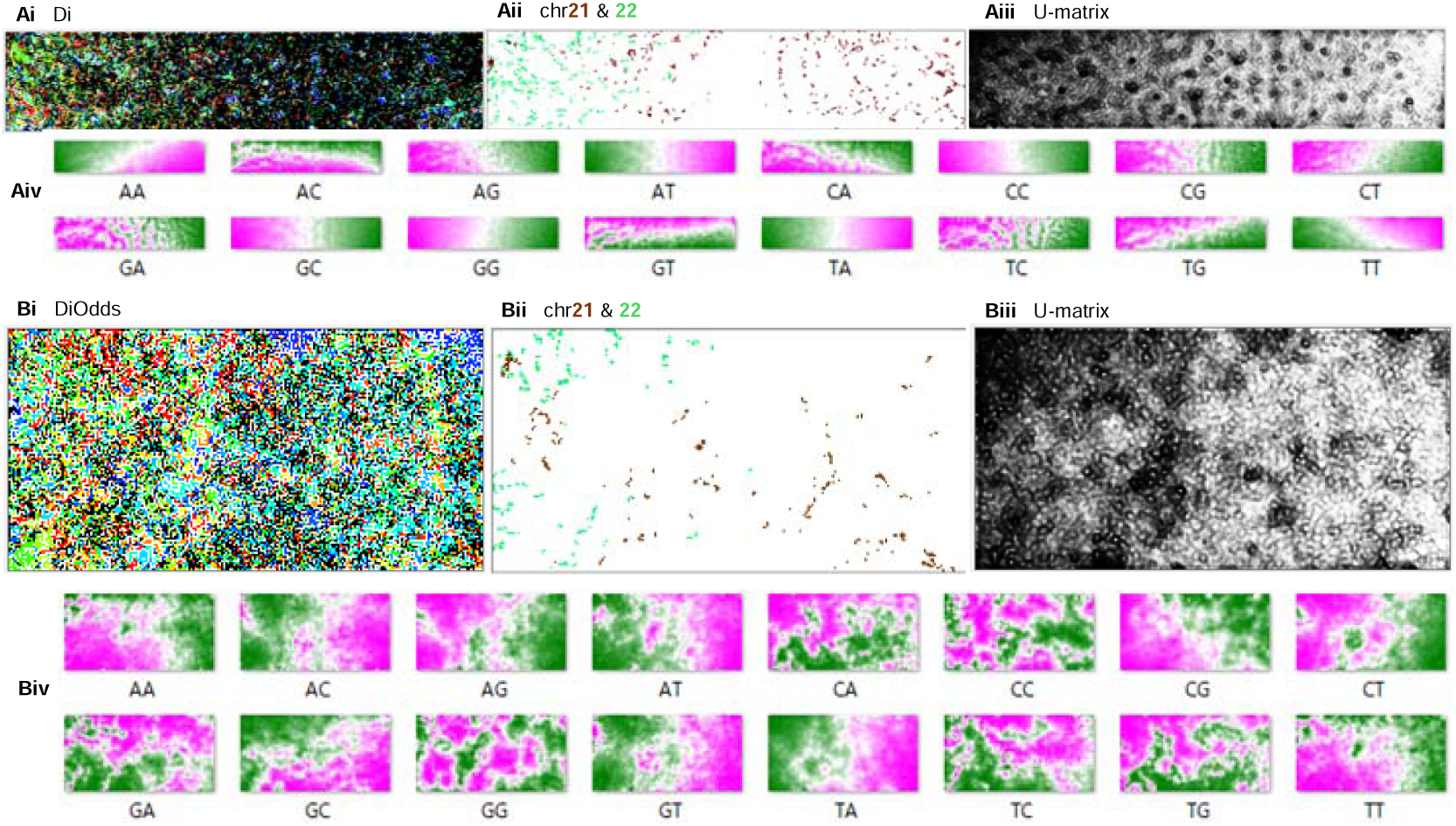
BLSOM of Di composition of 1-Mb windows across euchromatin. (**Ai**) BLSOM using standard Di composition. (Aii) Nodes containing chr21 or chr22 sequences are colored dark brown or light green, respectively. (**Aiii**) Node shading follows the method described in Fig. 1Bi. (**Aiv**) Heatmaps indicating the contribution level of each Oligo to clustering: pink (high), white (moderate), green (low). (Bi–Biv) present the BLSOM results for DiOdds, as in (Ai–Aiv).

A sliding□window approach is typically used to reduce segmentation effects and smooth local variation, but a 10□kb step is finer than required for these purposes. Because adjacent windows shifted by 10□kb have nearly identical Oligo compositions, the resulting map should display dotted, continuous trajectories. When the analysis encounters sharp composition shifts, such as band or isochore boundaries with evident G+C% changes, these trajectories may be interrupted. In these regions, BLSOM’s majority-vote learning assigns sequences spanning the boundaries to distant nodes representing compositionally stable domains, such as the main bodies of adjacent bands. Therefore, the BLSOM method should effectively reveal segmental structures of Oligo composition. We first examine the standard Di composition and then the odds ratio. In BLSOM of Di composition across euchromatin (Fig. 2Ai), most nodes are black, indicating limited chromosome□dependent separation. Figure 2Aii shows nodes corresponding to the two shortest chromosomes, chr21 (dark brown) and chr22 (light green). High□G+C% chr22 sequences (Dunham et al. 1999) map mainly to the left, whereas chr21 sequences, which include both low□ and high□G+C% regions (Hattori et al. 2000), span both sides. U-matrix (Fig. 2Aiii) reveals many small open circle-like zones delineated by clear black boundaries, although regions with faint or absent boundaries are also observed. As an explainable AI, BLSOM uses heatmaps (Fig. 2Aiv) to visualize contribution levels of individual Oligos to clustering at each node. Dis composed exclusively of A/T occur at high frequency (high contribution, pink) largely on the right, and at low frequency (low contribution, green) on the left: 1□Mb sequences with high A+T% cluster on the right, and those with high G+C% on the left, consistent with the chr21 and chr22 distributions in Fig. 2Aii.

The human genome is partitioned into large (>300□kb) G+C%□based segments known as isochores (Macaya et al. 1976; Bernardi et al. 1985). Consequently, analyses of Oligo composition, which is constrained by Mono composition, inevitably reflect this isochore structure. Several models have been proposed for isochore evolution (Bernardi and Bernardi 1986; Wolfe et al. 1989; Bernardi 1993), focusing on roles of mutation and selection. Mutation□driven effects are primarily reflected in Mono variation, whereas analysis of Oligo odds ratios (OligoOdds) highlights Oligo-sequence features independent of Mono bias, enabling more sensitive detection of selection□derived functional changes. BLSOM analysis of DiOdds (Fig. 2Bi) yields clearer patterns, with fewer black nodes than in Fig. 2Ai, showing greater chromosome-dependent separation. In Fig. 2Bii, chr22 sequences cluster predominantly on the left, whereas chr21 sequences appear on both sides, as shown in Fig. 2Aii. Notably, Fig. 2Bii contains fewer nodes than Fig. 2Aii, which indicates stronger chromosome-dependent clustering. The U□matrix reveals more numerous, circle□like zones than in Fig. 2Aii. The DiOdds heatmaps (Fig. 2Biv) also differ from the G+C%-based separation in Fig. 2Aiv. A/T Dis are not uniformly biased toward one side: AA and TT favor (pink) the left, whereas AT and TA favor the right. Likewise, CG and GC distributions are reversed between sides. Thus, OligoOdds BLSOM reveals clustering patterns that extend beyond simple G+C% segmentation.

### BLSOM Analysis of Tri to Hexa

BLSOM was extended to longer Oligos, and Tri- and Tetra-results are shown in Supplementary Fig. S2. In these cases, colored nodes progressively increase relative to Di. Here, we present the Penta case, which provides clearer results (Fig. 3A and B). Figure 3Ai shows the BLSOM pattern of the standard Penta, in which many nodes are marked in black; Fig. 3Aii highlights chr21 and chr22 nodes. The U-matrix distribution of small circle-like zones (Fig. 3Aiii) fairly resembles that in Fig. 2Aiii. With PentaOdds (Fig. 3Bi), however, colored nodes increase markedly and black nodes nearly disappear; chr21 and chr22 distributions (Fig. 3Bii) are simplified as observed for DiOdds (Fig. 2Bii). The U□matrix (Fig. 3Biii) further shows that most regions, particularly the A+T□rich right side, are covered by small circle zones clearly delineated by black boundaries reflecting distinct PentaOdds compositions.

**Figure 3.**
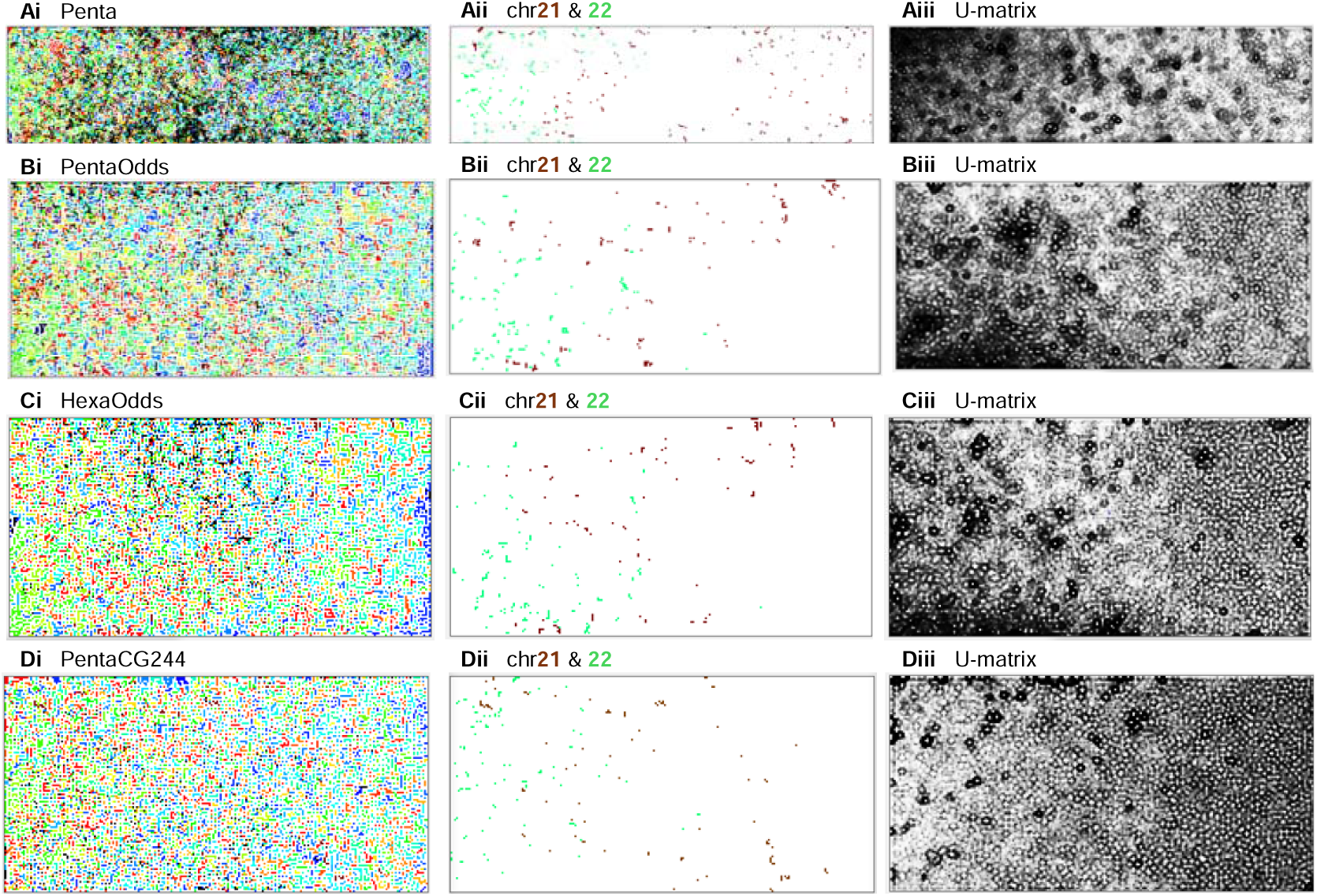
BLSOM for Penta and Hexa composition of 1-Mb windows in euchromatin. Results are shown for Penta (Ai), PentaOdds (Bi), HexaOdds (Ci), and PentaCG244 (Di). (Aii–Dii) Nodes containing chr21 or chr22 sequences are colored as in Fig. 2Aii. (Aiii–Diii) Node shading follows the method described in Fig. 1Bi.

HexaOdds results (Fig. 3Ci–iii) closely parallel PentaOdds, differing mainly in sharper boundaries in the U□matrix. Despite Hexa involving 4,096 variables versus 1024 for Penta, overall clustering structures are unexpectedly similar. Although ∼3,000 sequences are analyzed for each of chr21 and chr22, the number of nodes assigned is substantially smaller and similar between PentaOdds and HexaOdds (Fig. 3Bii, Cii). This shows that many 1-Mb sequences share similar PentaOdds/HexaOdds compositions and are consequently consolidated into single or adjacent nodes, which in turn implies Mb□scale compositional segmentation across euchromatin, as proven below. Collectively, analysis of OligoOdds, designed to probe functional features, enhances chromosome-dependent separation and reveals U□matrix patterns almost entirely covered by small circular zones. These findings highlight the unexpected capacity of unsupervised AI to uncover segmental genomic features, motivating further investigation into the biological basis of this segmentation.

### BLSOM Analysis of CG-containing Oligos

If the unexpectedly found ∼2,000 small circular zones reflect biologically meaningful genomic segmentation, similar patterns should also emerge with other Oligo types. To test this, we examine CG-containing Oligos, which have well-established roles in epigenomic regulation (Jones 2012; Smith et al. 2025). Figure 3D shows the BLSOM of 244 CG-containing Pentas (PentaCG244), focused on their relative frequencies rather than odds ratios. Compared with the non-odds Penta BLSOM in Fig. 3Ai, the number of variables is reduced to less than one-quarter, yet colored nodes increase markedly (Fig. 3Di): most nodes comprise sequences from a single chromosome, as observed for PentaOdds and HexaOdds (Fig. 3B, C). Despite clear differences in chr21 and chr22 positioning in Fig. 3Dii from the positioning observed for PentaOdds (Fig. 3Bii) and HexaOdds (Fig. 3Cii), the U□matrix is again filled with small circles (Fig. 3Diii). This supports the pivotal role of CG□containing Oligos in Mb□scale segmentation. Heatmaps explaining the contribution levels of CG-containing Pentas are presented in Supplementary Fig. S3A, along with their interpretations. The BLSOM of CG-containing Hexas (HexaCG1185) is shown in Supplementary Fig. S3B, which displays a pattern closely matching PentaCG244 (Fig. 3D). To highlight functional features, we analyzed OligoOdds and CG□containing Oligos and found highly similar U-matrix partitions across Oligos of varying lengths and types, indicating that the underlying segmental structures reflect fundamental functional divisions within the genome.

### Similar Oligo-Usage Zones

Detailed examination of small circle zones in the U-matrix revealed occasional morphological variations, including rod-like, curved rod-like, and gourd-like forms. The U-matrices of PentaOdds, HexaOdds, PentaCG244, and HexaCG1185 contained over 1,800 sharply defined zones, significantly exceeding 1,900 when faint boundaries were considered. These are hereafter referred to collectively as ∼2,000 small-circle zones, and the similarity of these four zonings will be confirmed by different approaches later. Adjacent zones are separated by black lines, denoting distinct Oligo compositions, whereas nodes within each zone are nearly white, indicating high internal similarity. Because nearly the entire BLSOM is dominated by colored nodes, sequences of individual nodes originate from a single chromosome. To compute Oligo composition, we analyzed ∼280,000 overlapping 1-Mb sequences generated with a 10-kb increment. Because adjacent windows differ by only 10 kb, they yield largely redundant Oligo compositions, and neighboring sequences are expected to fall largely within the similar Oligo-usage zones except near segmentation boundaries. For example, chr21 yields 3,290 sequences, yet only ∼50 brown nodes appear in Fig. 3Bii–Dii, indicating clear clustering of many sequences into the same or neighboring nodes. However, under a majority-rule-based machine-learning framework, sequences spanning segmentation boundaries are expected to be assigned to distant nodes occupied by sequences that are derived from the main body of the segment on either side, creating a characteristic “tearful separation.” To test this expectation, we analyzed the PentaOdds BLSOM in Fig. 3B, tracking the horizontal (X) and vertical (Y) coordinates of the nodes assigned to each 1□Mb fragment of chr21 (i.e., dark brown nodes in Fig. 3Bii).

The BLSOM comprises 27,664 nodes (247□×□112), and Fig. 4A displays the X/Y-coordinate values of the ∼50 brown nodes along chr21 euchromatin for PentaOdds BLSOM (Fig. 3Bii). Sequences within each small-circle zone in the U-matrix share identical or nearly identical coordinates, resulting in horizontal lines with minimal X/Y variation. At sharp transitions in oligo composition, sequences flanking the transition boundary map to distinct node zones, yielding abrupt shifts in their X/Y coordinates. Because the initial BLSOM configuration was defined by the first and second principal components, fluctuations along the X and Y axes capture variation driven by different underlying factors. As shown in Fig. 4A, both axes fluctuate in broadly similar positions but in opposite directions, producing highly jagged X and Y profiles. Green triangles mark 850-band boundaries, yet abrupt changes in both axes occur at many additional positions. Figure 4B plots chr21 X and Y data from HexaOdds BLSOM (Fig. 3Cii), and Figs. 4C and D from PentaCG244 and HexaCG1185 BLSOMs (Fig. 3Di and Supplementary Fig. S3Bii).

**Figure 4.**
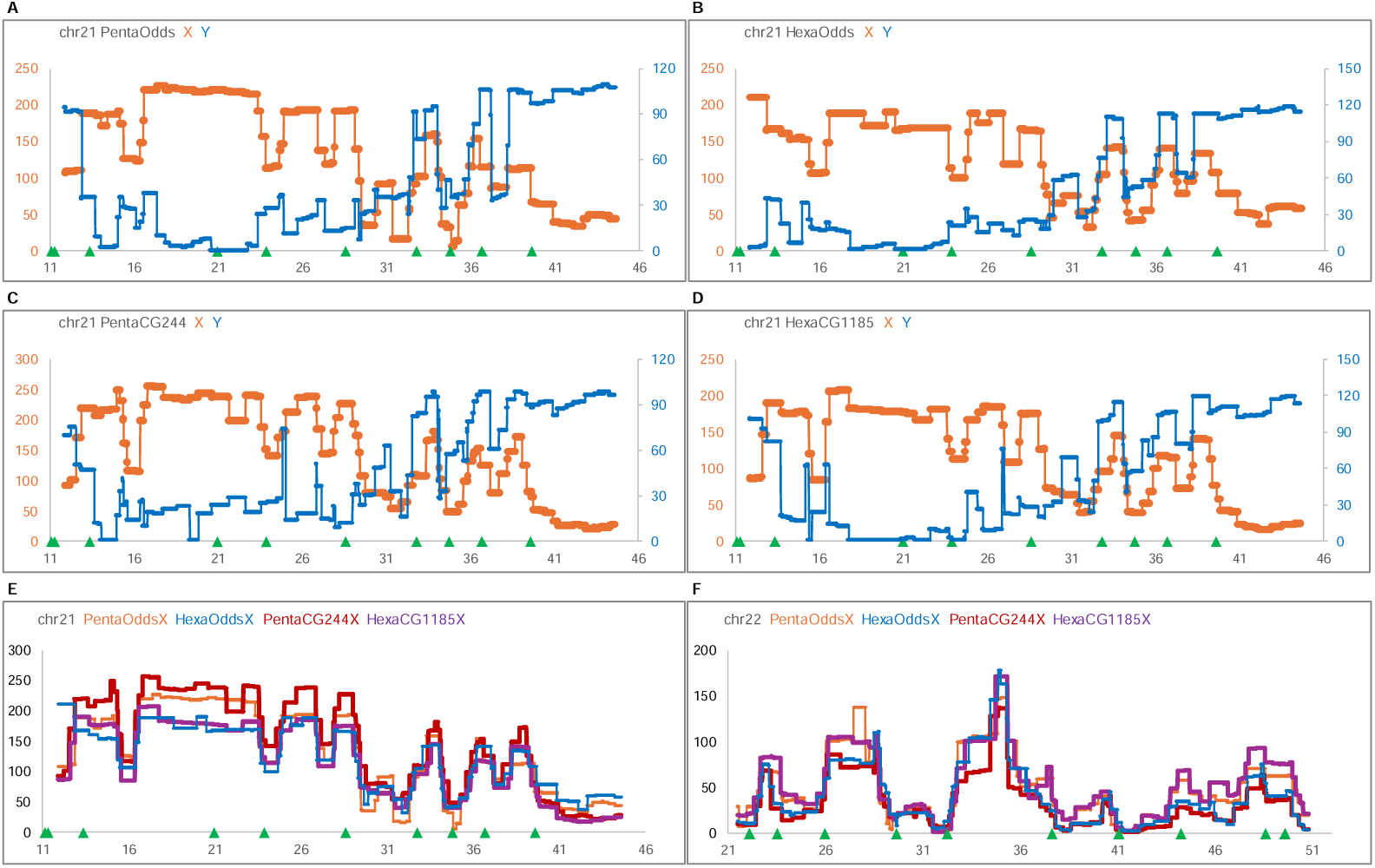
Variation of BLSOM X- and Y-axis values along chr21 euchromatin. X-values (brown) and Y-values (blue) are plotted along chr21 for PentaOdds (**A**), HexaOdds (**B**), PenCG244 (**C**), and HexaCG1185 (**D**). Numbers below the horizontal axis indicate genomic coordinates (Mb), and green triangles denote the positions of the 850LJband boundaries in this and subsequent figures. (**E, F**) X-values of these four BLSOMs are plotted along chr21 and chr22, respectively. Oligo type and color correspondence is shown at the top of the figure.

In Figs. 4E and 4F, the X-values of these four BLSOMs are plotted along chr21 and chr22 euchromatin. Notably, HexaOdds BLSOM uses 4,096 variables, approximately 17 times more than PentaCG244 BLSOM, and these focus on two distinct features: the OligoOdds and the relative abundance of CG-containing Oligos. Despite this clear difference, it produces comparable fluctuating profiles, which is consistent with the similar U-matrix patterns that delineate ∼2,000 circular zones. The shared jagged profiles in Fig. 4 therefore likely reflect the genome’s fundamental segmented architecture. Genomic regions in which the X and Y values show minimal variation have similar Oligo compositions and are referred to as similar Oligo-usage zones, which likely correspond to the small-circle zones in the U-matrix. Table 1 summarizes correlations of variation along the X- and Y-axes among four Oligo types for chr21 and chr22, showing high coefficients for either axis or chromosome. In PentaCG244 and HexaCG1185, sharp slender Y-axis peaks are observed at X-axis fluctuation sites (Fig. 4C, D). The reason why the correlation coefficient is lower on the Y-axis will be discussed later in connection with these slender peaks and local structural features at band boundary regions.

**Table 1.**
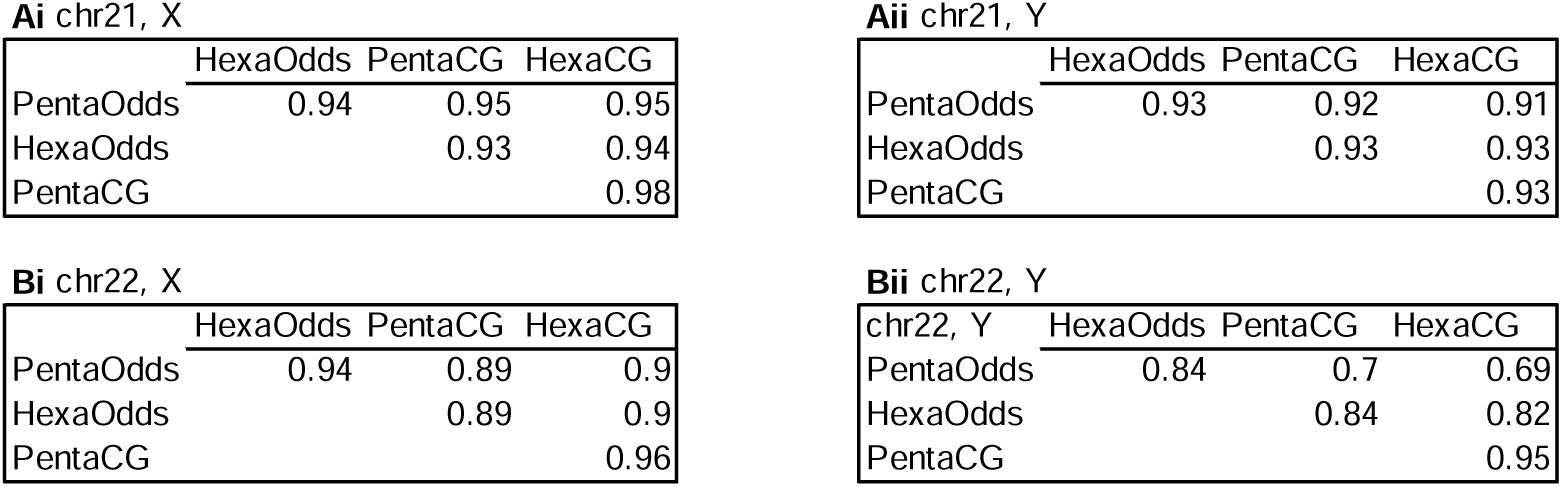
Pearson correlation coefficients for variation along the X and Y axes among four Oligo types. The numbers of data points for chr21 and chr22 are 3,290 and 2,952, respectively.

### Strategies for Band Reconstruction using Non-AI Informatics

We next examine how Mb-scale similar Oligo-usage zones identified by BLSOM correspond to chromosome bands, using band positional data and non-AI informatics. The UCSC Genome Browser defines band boundaries at single□base resolution based on FISH data of 850 bands. However, these 850 bands likely encompass finer bands detectable at the 2000□band resolution (Yunis 1981). The U-matrix reveals ∼2,000 similar Oligo-usage zones, making the 2000-band pattern a suitable reference for comparing chromosome bands to these zones. However, positional data for 2000 bands are unavailable. We thus use the 850-band data to assess Oligo-usage differences between Giemsa-positive and -negative bands and to test the following two hypotheses for informatics-based band reconstruction. In the UCSC Genome Browser, Giemsa-stained bands of euchromatin are annotated as gneg or gpos100, with intermediate categories gpos25, gpos50, and gpos75, representing increasing levels of Giemsa staining intensity. We focused on gneg and gpos100, the most distinct classes, and extracted all sequences assigned to these two categories. These sequences cover 63% of euchromatin and form the basis of this band-reconstruction analysis; intermediate categories are addressed separately below.

Hypothesis 1: While the finer 2000 bands are generally embedded within the broader 850 bands, extracting all sequences classified as gneg or gpos100 using the 850-band coordinates should yield diagnostic Oligo sets distinguishing Giemsa-positive and -negative bands. Such sets may enable informatics-based reconstruction of pseudo-bands.

Hypothesis 2: Diverse biological functions occur mainly during interphase rather than metaphase. If bands truly reflect functional segmentation, then diagnostic Oligo sets that accurately capture this functional segmentation would reconstruct the finer 2000 bands observed in prophase, which is closer to interphase.

### Investigating the Diagnostic OligoOdds Set

To identify diagnostic Oligos distinguishing gpos100 and gneg, we extracted all euchromatin sequences corresponding to these categories using UCSC coordinates and computed their OligoOdds. The OligoOdds for the entire euchromatin was also calculated as a normalization reference. For each Oligo, OligoOdds values for two band types are normalized against the euchromatin reference, and these normalized data are expected to yield diagnostic Oligos that predict band types and support functional interpretation of their differences. Figure 5Ai shows normalized DiOdds for all 16 Dis. In gpos100, TA is highest (red background), whereas the next two ranks are not A/T-only and the fifth is GC, showing that these preferences are not attributable solely to A+T richness of gpos100. The least favored Dis is CG (green letter), followed by CT, AG, AA, and TT; except for CG, these are not C/G-only. In gneg, the preference is reversed, with AA and TT ranking fourth and fifth, showing that these preferences cannot be explained simply by G+C richness of gneg. As mentioned later, consecutive A or T is more strongly favored in gneg when longer OligoOdds are analyzed. The gpos100/gneg (dark red) is presented as an indicator of G-positivity, reflecting the extent to which a Di is favored in gpos100 and disfavored in gneg. Supplementary Fig. S4 shows results of three intermediate bands; gpos75 resembles gpos100, gpos25 resembles gneg, and gpos50 is intermediate. For clarity, the following analyses focus on gneg and gpos100, while features of other three bands are described later.

**Figure 5.**
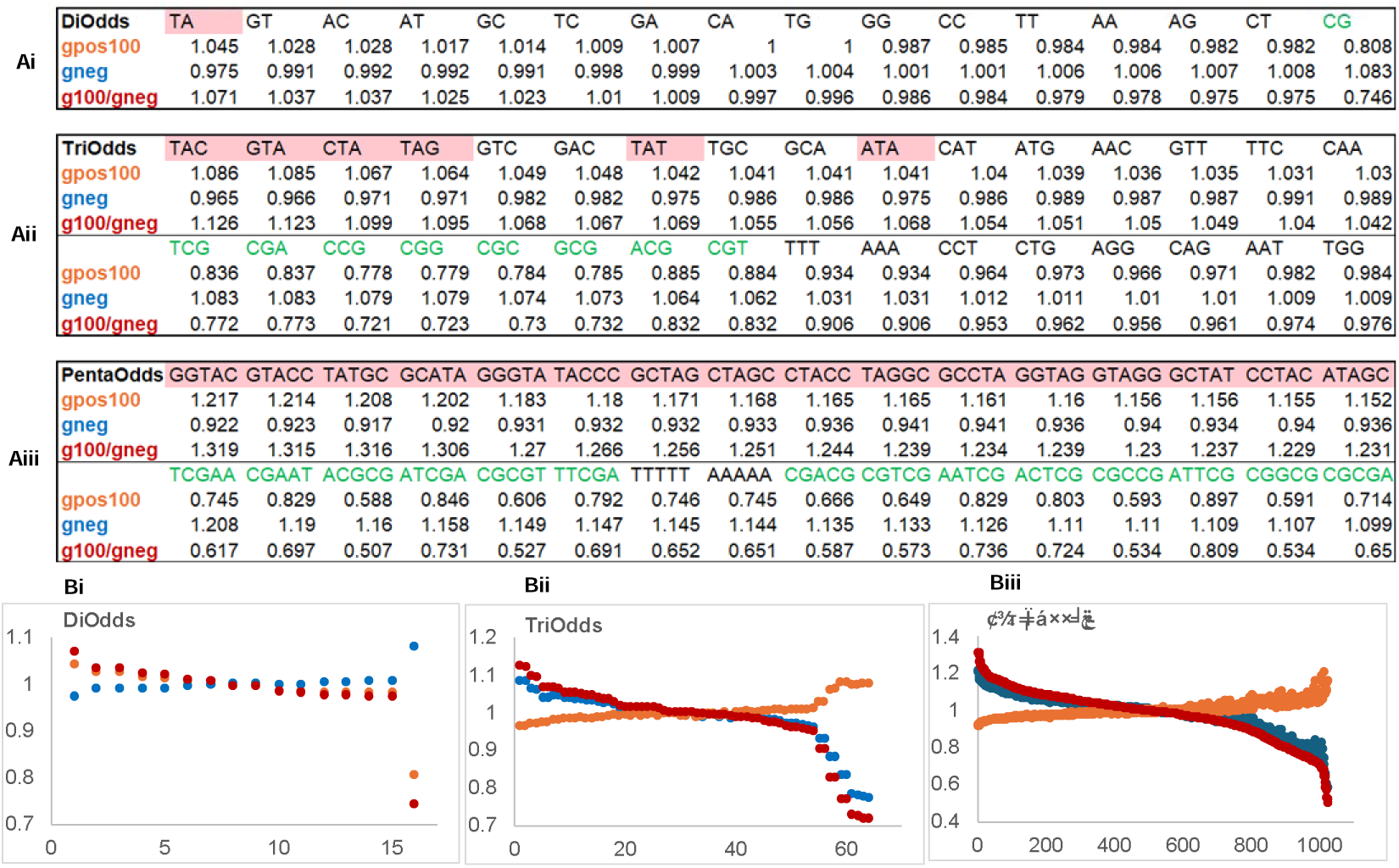
Identification of diagnostic Oligos. (**Ai**) Normalized DiOdds values for gpos100 and gneg, and their ratios (g100/gneg, dark red), are ranked by the ratio; TA and CG are highlighted (red background and green letter). (**Aii, Aiii**) Top 16 normalized TriOdds and PentaOdds enriched in gpos100 and gneg are shown above and below the horizontal line, with TA- and CG-containing Oligos highlighted as in (**Ai**). (**Bi–Biii**) Normalized DiOdds, TriOdds, and PentaOdds for gpos100, gneg, and their ratio (g100/gneg) are shown in a scatter plot ranked by their ratios. Symbol coloring follows (Ai–Aiii).

Figure 5Aii presents the top 16 Tris favored in gpos100 (above the horizontal line) or gneg (below). In gpos100, TA-containing Tris are strongly preferred, particularly when flanked by C/G, indicating preferences beyond simple A+T richness. In gneg, all eight CG-containing Tris rank highest, with A/T flanks leading, followed by AAA and TTT; GC-containing Tris other than CG-containing ones do not rank highly, showing that gneg order reflects sequence features rather than G+C richness. Extension of the analysis to PentaOdds confirmed these trends; Figure 5Aiii shows that gpos100 favors TA-containing Pentas, especially when flanked by C/G. In gneg, CG-containing Pentas dominate, particularly with A/T flanks, though distinct CGCG-containing sequences also appear among the top, suggesting a more complex pattern. Notably, AAAAA and TTTTT rank seventh and eighth.

We next consider biological relevance of diagnostic OligoOdds. Segmental mutation pressures (Wolfe et al. 1989) have shaped large-scale genomic structures, isochores. Functionally important Oligo sites tend to resist these pressures and thus become more distinct. OligoOdds analysis therefore provides an efficient approach for identifying Oligo sequences with functional significance. TA is known to be highly flexible and bendable, facilitating stable nucleosome formation (Trifonov and Sussman 1980; Takasuka and Stein 2010), consistent with gpos100 representing dense chromatin. In contrast, consecutive A/T sequences (e.g., AAAAA/TTTTT) tend to impede chromatin formation (Segal and Widom 2009) and are enriched in the more open chromatin of gneg. Cytosine in CG is the methylation site, and neighboring bases modulate the binding of epigenetic factors such as methyl□CpG–binding proteins, highlighting the functional diversification of CG□containing Oligos (Smith et al. 2025; Deaton and Bird 2011). Figure 5Biii illustrates the normalized gpos100 and gneg PentaOdds values, along with their ratio representing a G□positivity indicator. The dark-red curve of this indicator reveals that ∼5% of Pentas at both extremes deviate sharply from the rest, as indicated by steep slope (Fig. 5Biii). We therefore select the corresponding top and bottom 51 PentaOdds as a diagnostic oligo set for gpos100 and gneg, respectively, and use them for informatics-based band reconstruction. The 102 focal Pentas are shown in Supplementary Fig. S5.

### Indices of G-positivity and Their Variation Along Chromosomes

Averaging the 51 diagnostic PentaOdds ratios (g100/gneg in Fig. 5) for gpos100 and gneg produced two band-specific indices, whose variations across chr21 euchromatin are shown in Fig. 6Ai. The gpos100 and gneg profiles (dark and light green) are approximately mirror□symmetric and strongly anticorrelated (r□=□-0.94), confirming that these indices capture band-specific features. For direct comparison with segmental structures revealed by other methods, a single G-positivity index is more practical than two separate indices. We therefore calculated the ratio of two indices as a unified G-positivity index for every 1-Mb sequence. Its variation along chr21 is shown in dark red in Fig. 6Aii. Bands are known to correlate with G+C% (and A+T%) variation, and a comparison of the G-positivity index with Mb-level A+T% variation (blue) reveals similar profiles (r = 0.94), although the G-positivity index is derived from a odds-ratio analysis that is free from Mono bias. Elevated G-positivity corresponds to high A+T%, whereas a reduced value corresponds to low A+T%. Notably, G-positivity fluctuates more sharply than A+T%, highlighting its diagnostic utility for band reconstruction.

**Figure 6.**
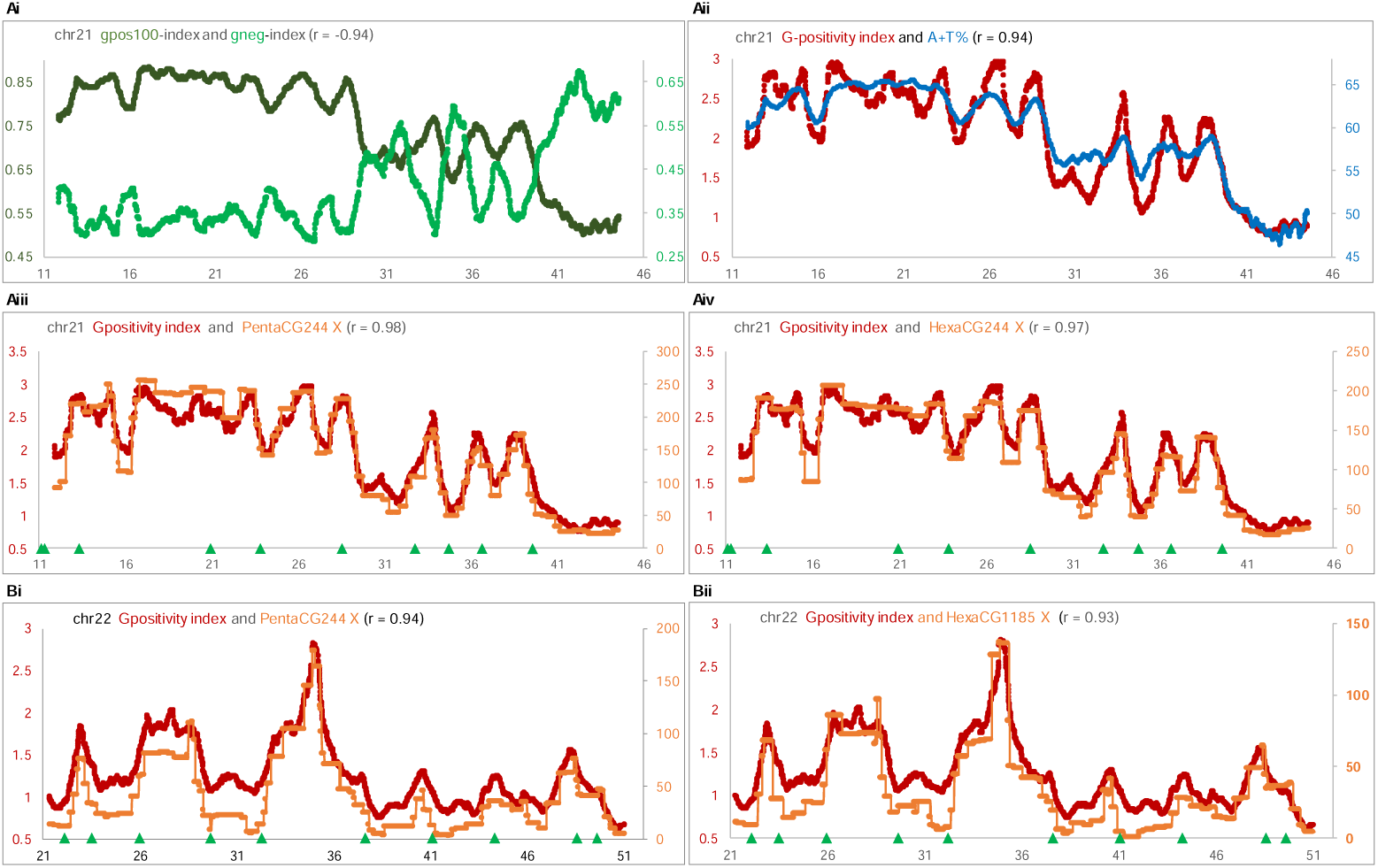
Variation of G-positivity along chr21 and chr22 euchromatin. (**Ai**) Mean of 51 PentaOdds values for gpos100 and gneg diagnostic sets. The gpos100- and gneg-indices (dark and light green) are plotted along chr21. (**Aii**) G-positivity index (dark red) and A+T% (blue) plotted along chr21. (**Aiii, Aiv**) G-positivity index and BLSOM X-values for PentaCG244 (**Aiii**) and HexaCG1185 (**Aiv**) are plotted along chr21 in dark red and brown. (**Bi, Bii**) G-positivity index and BLSOM X-values for PentaCG244 (**Bi**) and HexaCG21185 (**Bii**) are plotted along chr22, using the same color scheme as in (**Aiii, Aiv**).

We next analyzed how changes in the G-positivity index relate to shifts in BLSOM coordinates along chr21 euchromatin. In Fig. 6Aiii and Aiv, the G-positivity profile (dark red) closely follows the X-axis variation in the PentaCG244 and HexaCG1185 BLSOMs (light brown in Fig. 4Ai and Supplementary Fig. S3), with correlation coefficients of 0.98 and 0.98. Similar results were obtained for chr22 (Fig. 6Bi, Bii). Beyond expectation, the G-positivity profile obtained with non-AI informatics and 850-band coordinates closely recapitulates the segmental structures detected by BLSOM on chr21 and chr22. The positions and orientations of the transitions are consistent, although the BLSOM transitions are sharper than the G-positivity transitions. This strong concordance indicates that both approaches capture a common underlying segmentation of biological relevance, likely reflecting to the 2000 prophase Giemsa bands, thereby supporting Hypothesis 2.

### Mapping of Sequences from Five Band Types and Their Boundaries onto PentaOdds BLSOM

To reconstruct bands, we defined a G-positivity index based solely on gpos100 and gneg features. In Supplementary Fig. S4A, DiOdds showed intermediate characteristics for remaining three-band types, whereas higher□order features captured by PentaOdds may provide clearer discrimination among band types. Next, we analyze locational characteristics of sequences of all five-bands on the PentaOdds BLSOM in Fig. 3Bi. Band sequences of each type were extracted from 850 bands, and PentaOdds values were computed using a 1-Mb window shifted every 10□kb. Because terminal regions cannot be fully shifted, sub-Mb gaps unavoidably emerge. The 1-Mb gap is negligible at the chromosome scale but becomes substantial at the band scale (mean length ∼3.5□Mb). To mitigate this, residual sub-Mb sequences were further shifted in 10-kb steps to 250 or 500□kb, and the resulting counts were included. Since future studies are expected to address band boundaries, which may contain distinctive Oligo-composition patterns as discussed later, we included boundary sequences. For each boundary, a 1-Mb region spanning 500 kb upstream and downstream was examined. Since this 1-Mb sequence provides only one data point, we again incorporated additional data down to 250 or 500 kb. Although 1-Mb sequences obtained from 850 bands differ from those used to construct the original BLSOM, the 10-kb step produces highly similar sequences that map to corresponding nodes. However, Oligo frequencies of sub-Mb fragments may reveal local structures invisible at the Mb scale, enabling analysis of boundary□specific features. We applied 250□ and 500□kb cutoffs to identify the window size at which such local structures become detectable.

**Figure 7.**
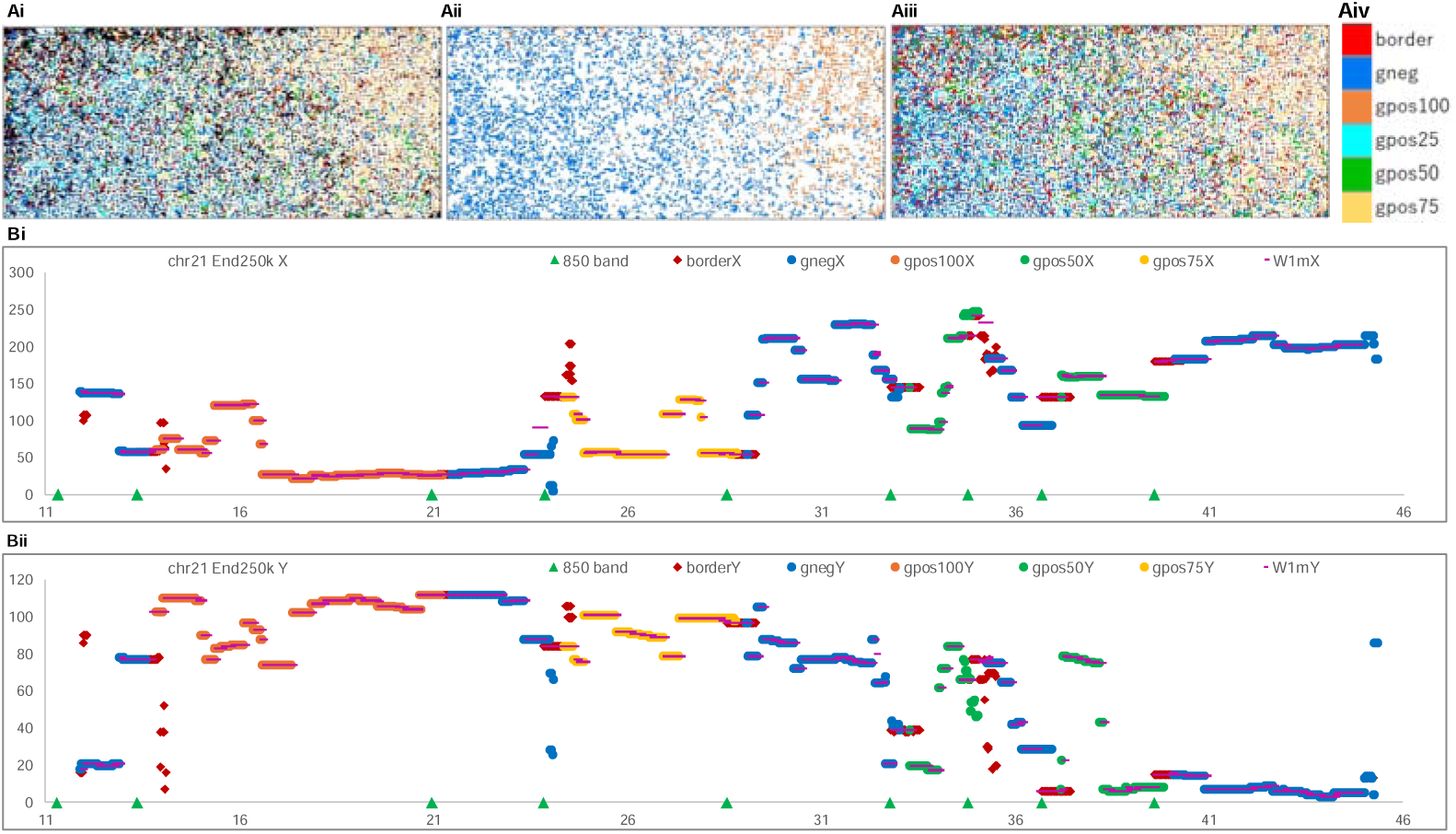
Mapping of PentaOdds of sequences from five bands and their boundaries onto the PentaOdds BLSOM. (**Ai**) PentaOdds data for five bands and their boundaries, including residual segments down to 250 kb, are mapped onto the PentaOdds BLSOM in Fig. 3Bi. Nodes containing sequences from a single category are color-coded, whereas mixed nodes are black. (**Aii**) Nodes containing gpos100 or gneg sequences are color-coded. (**Aiii**) Nodes colored by predominant category. (**Aiv**) Category and color correspondence. (**Bi, Bii**) Variation in the XLJ and Y-values (**Bi and Bii**) of the BLSOM in panel (**Ai**) along chr21. Colored symbols representing the six categories are shown at the top of each panel. W1mX/Y (pink) denotes X/Y axis values of the PentaOdds BLSOM (Fig. 3Bi) constructed using only 1-Mb sequences.

In Fig. 7Ai, PentaOdds data of 1-Mb fragments from five-band types and their boundaries, including residual segments down to 250 kb, were mapped onto the PentaOdds BLSOM in Fig. 3Bi, by assigning each fragment to the node with the closest composition. Nodes containing one category are color-coded, whereas mixed nodes are shown in black. In Fig. 7Aii, nodes containing only gpos100 or gneg sequences are color□coded; gpos100 sequences (royal blue) are enriched on the left, while gneg sequences (mandarin orange) are enriched on the right, producing a pattern that broadly reflects the band types. In Fig. 7Aiii, each node is colored by its predominant category. Differences among gpos25, gpos50, and gpos75 sequences generally align with the DiOdds features described in Supplementary Fig. S4A, while reveal additional characteristics. Gpos25 (cyan) resembles gneg in its right-side abundance, yet it forms distinct small territories. Gpos75 (khaki) resembles gpos100 in its left-side enrichment. Gpos50 (lime green) is more broadly distributed; it also forms unique territories mainly in the central region, indicating sequence groups specific to gpos50. Boundary sequences (red) are dispersed across the map without forming distinct territories, though frequently appear in black nodes in the upper left and along the left edge of Fig. 7Ai. The original BLSOM used for mapping provides finer U-matrix partitions approaching the 2000-band resolution, whereas mapped sequences are color-coded under the 850-band scheme. Broad gneg bands generally encompass narrow sub-bands observed under the 2000-band scheme, such as gpos100-like narrow bands. Although labeled as gneg (royal blue), these narrow segments should map to the gpos100 territories enriched in mandarin orange nodes, and many such cases are evident in Fig. 7Aii. This increases visual complexity of Fig. 7A, yet the genomic coordinates of these narrow segments help predict banding patterns beyond the 850-band resolution.

### Possible Local Structures at Band Boundaries

X- and Y-axis values for band and boundary sequences of chr21 in Fig. 7Ai are plotted in Fig. 7Bi and Bii. This BLSOM corresponds to the original map in Fig. 3B, and the axis values for the original 1-Mb sequences are indicated by thin pink lines. Sub-Mb sequences of bands, particularly those at band boundaries (dark red), often show pronounced shifts from the pink line. When local structural features exist at boundaries, they appear as clear upward or downward shifts rather than intermediate values between adjacent bands. These shifts arise because the original BLSOM was constructed only from 1-Mb sequences, which lacks nodes capturing compositional features detectable only at ∼250□kb, thereby forcing sub-Mb sequences to map to the nearest compositionally similar nodes. Consequently, their mapped positions lack biological meaning, simply showing the presence of local structures. Analyses using 500-kb terminal extensions (Supplementary Fig. S6) detected fewer local structures than those using 250□kb, indicating that a 250-kb window improves detection of boundary-specific features.

## DISCUSSION

### Exploration of Band Boundaries and Future Perspectives

Unsupervised AI can uncover unexpected insights. The BLSOM analysis of 1-Mb moving windows with a 10-kb step identified ∼2,000 zones with similar Oligo usage. Although the primary goal was to examine their correspondence to band structures, preliminary information on band boundaries also emerged (Fig. 7). Boundaries represent transitions between adjacent bands; when features of both bands fade gradually, precise boundary localization is difficult and of limited biological relevance. By contrast, when band structures reflect functional segmentation, boundaries are expected to exhibit distinctive local features, indicating that accurate boundary estimation and functional interpretation are feasible, thereby opening new avenues for research.

Because the initial BLSOM configuration was defined by the first and second principal components, fluctuations along the X and Y axes capture variation driven by different underlying factors. In this study, X-axis variation has been the primary focus, whereas the Y□axis has received far less attention. Although the biological basis of Y-axis variation remains largely unexplored, sharp and rod-like peaks appear at positions exhibiting X-axis shifts primarily for PentaCG244 and HexaCG1185 (Fig. 4C, D). Focusing on CG-containing Oligos should reveal boundary features detectable even at the Mb scale. As shown in Table 1, the coefficients of variation for the four Oligo types on both chr21 and chr22 are lower along the Y axis than the X axis. This indicates that fluctuations along the Y axis considerably reflect variation dependent on Oligo types, and factors driving Y-axis variation may offer further insight into functional genome segmentation. Characterizing band-boundary features would enable sequence-based inference of band architecture as functional units across primates and potentially other mammals, thereby providing a basis for understanding band-boundary function and helping to bridge classical cytogenetics with modern, AI-driven genomic analyses.

### Additional Findings

Our primary focus was to assess the biological relevance of AI-identified similar Oligo-usage zones, but we also observed additional findings related to differences among chromosomes. ChrX displays features distinct from autosomes, including distinct blue zones in Fig. 2Bi and 3Ai–Ci, and dense-boundary circles in the U-matrix (Fig. 2Biii and 3Aiii–Ciii) align well with these blue chrX zones. Oligo-composition analysis reveals elevated frequencies of CC/GG-containing motifs across multiple chrX-specific regions. This is evident in the pseudoautosomal region (PAR), which pairs with chrY during meiosis. Given the established association between C/G-tract motifs and recombination mechanisms (Saranathan et al. 2019; Camarillo et al. 2021), this enrichment on sex chromosomes is noteworthy.

Analyses of diagnostic OligoOdds can show band□boundary characteristics, and TriOdds profiling reveals strong enrichment of oligopurine/oligopyrimidine motifs, such as AGG, AAA, CCT, TTT, and GAG (Supplementary Fig. S4C). Because such oligopurine/oligopyrimidine motifs readily form non□B structures, including DNA-only and RNA□associated triplexes (Bacolla et al. 2015), they may underlie band boundary structures and nuclear compartmentalization (Ohno et al. 2002; Wang et al. 2023; Makova and Weissensteiner 2023).

## METHODS

### Human Genomic Sequences

The telomere-to-telomere (T2T CHM13v2.0) human genome assembly (Nurk et al. 2022) was obtained from the UCSC Genome Browser. Centromeric and pericentromeric constitutive heterochromatin, which is enriched in diverse repetitive elements and shows characteristic BLSOM clustering in Fig. 1, was excluded from the analysis of euchromatin, as described below. Oligo compositions of centromeric constitutive heterochromatins in GRCh38 were characterized previously (Wada et al. 2020; Iwasaki et al. 2022). Their sequence coordinates were converted to the T2T assembly using LiftOver (https://github.com/burgshrimps/liftover_T2T) and were then used to extract euchromatic sequences (Supplementary Table). T2T sequence coordinates for bands assigned to eight band types categorized in the UCSC Genome Browser were used to extract the bands and their boundary sequences.

## BLSOM

The Self-Organizing Map (SOM) is an unsupervised neural-network algorithm for clustering and visualizing high-dimensional data (Kohonen 1990). We adapted SOM for genome informatics using batch learning, making the learning process and resulting map independent of input order (Kanaya et□al. 2001). Initial weights were set by principal component analysis, and weight vectors were arranged on a two□dimensional lattice and updated as previously described (Abe et al. 2003). The lattice spanned five standard deviations of the first two principal components. During learning, each genomic fragment was assigned to the node with the smallest Euclidean distance in Oligo-composition space, and node weight vectors were iteratively updated as described in detail (Abe et al. 2003): see also Supplementary Method. Oligo contributions at each node were visualized with a pink-to-green heatmap, representing high to low contributions (Kanaya et al. 2001). A Python implementation of BLSOM is available at https://doi.org/10.6084/m9.figshare.25036358.v1, and a Linux version is provided at http://bioinfo.ie.niigata-u.ac.jp/?BLSOM.

## ACKNOWLEDGMENTS

This work was supported by research grants from the Japan Science and Technology Agency (JST CREST Grant Number JPMJCR20H1; To T.A.). Open access funding was provided by Nagahama Institute of Bio-Science and Technology.

## Contributions

T.I. performed the data analysis, oversaw the project, and wrote the manuscript. K.W. was responsible for programming. Y.I. and Y.W. validated the software. T.A. designed the project and obtained the grant.

## Data availability

All data generated or analyzed during this study are included in this published article and its Supplementary files.

## Code availability

A Python implementation of BLSOM is available at https://doi.org/10.6084/m9.figshare.25036358.v1, and a Linux version is provided at http://bioinfo.ie.niigata-u.ac.jp/?BLSOM.

## Supplemental Material

This file includes:

Supplementary Figures S1 to S6

Supplementary Table

Supplementary Method

